# Towards Digital Quantification of Ploidy from Pan-Cancer Digital Pathology Slides using Deep Learning

**DOI:** 10.1101/2024.08.19.608555

**Authors:** Francisco Carrillo-Perez, Eric M. Cramer, Marija Pizurica, Noemi Andor, Olivier Gevaert

## Abstract

Abnormal DNA ploidy, found in numerous cancers, is increasingly being recognized as a contributor in driving chromosomal instability, genome evolution, and the heterogeneity that fuels cancer cell progression. Furthermore, it has been linked with poor prognosis of cancer patients. While next-generation sequencing can be used to approximate tumor ploidy, it has a high error rate for near-euploid states, a high cost and is time consuming, motivating alternative rapid quantification methods. We introduce PloiViT, a transformer-based model for tumor ploidy quantification that outperforms traditional machine learning models, enabling rapid and cost-effective quantification directly from pathology slides. We trained PloiViT on a dataset of fifteen cancer types from The Cancer Genome Atlas and validated its performance in multiple independent cohorts. Additionally, we explored the impact of self-supervised feature extraction on performance. PloiViT, using self-supervised features, achieved the lowest prediction error in multiple independent cohorts, exhibiting better generalization capabilities. Our findings demonstrate that PloiViT predicts higher ploidy values in aggressive cancer groups and patients with specific mutations, validating PloiViT potential as complementary for ploidy assessment to next-generation sequencing data. To further promote its use, we release our models as a user-friendly inference application and a Python package for easy adoption and use.

## Introduction

Ploidy refers to the number of sets of chromosomes in a cell. Normal human cells have two sets (diploid). In contrast, cancer cells often exhibit an abnormal number of chromosomes, referred to as aneuploidy (as result of insertion or deletion). Ploidy can be estimated by measuring the amount of DNA in a cell, referred to as DNA ploidy value.^1^. This abnormal number of chromosomes has been linked with more aggressive tumors, poorer prognosis, and a higher chance of metastasis in some cancer types^2–4^.

Several techniques exist to quantify ploidy in biology. Traditional karyotyping allows direct visualization of chromosomes but is labor-intensive, limited to a small number of cells and lacks sensitivity for smaller changes^5^. Next-Generation Sequencing (NGS) excels in sensitivity for various genomic alterations, but comes with higher costs, complex data analysis^6^, and error-prone ploidy quantification for near-euploid states. Two other techniques are mainly used nowadays to quantify ploidy from tissue samples: flow cytometry^7,8^ and image-based analysis using immunofluorescence (IF)^9,10^. However, they both present drawbacks. Flow cytometry is confounded by cell cycle state. For example if cells tend to spend longer in G2M for any reason it will look like an increase in ploidy, when in fact only a shift in cell cycle state duration has happened. On the other hand, with IF imaging, it can be difficult to obtain unstained tissue sections, and sampling errors may arise from manual counting in a few histological regions.

During clinical practice, one of the main cancer diagnosis tools used by pathologists are tissue samples stained with hematoxilyn & eosin (H&E). These tissue samples are routinely obtained and digitized as whole-slide images (WSI) (also known as digital pathology slides) for visual examination^11,12^. Thus, having a tool that quantifies ploidy from WSI samples can result in a fast and cheap quantification tool that could be integrated into the clinical practice, without the requirement of processing a tissue sample.

Machine learning (ML), particularly deep learning, has achieved remarkable results in digital pathology, enabling various cancer-related tasks^13–18^. Traditionally, models pre-trained on natural images were fine-tuned for specific WSI tasks. However, the success of self-supervised learning (SSL) has boosted the development of WSI-specific SSL models^19–21^. These models reveal a unique latent space within WSIs, offering the potential to significantly enhance downstream task performance by using the model output’s features. These novel feature extractors are usually based on transformer-based models, specifically vision transformers, that are an extension of natural language processing transformers applied to imaging modalities^22^. The improvements achieved by using there feature extractors over using Imagenet-trained models is task-specific, and needs to be tested for each independent task.

Regarding the development of ploidy quantification models from H&E images, previous work has resulted in a deep learning-based model to quantify hepatic ploidy from non-cancerous H&E images with good results^23^. However, authors did not predict the bulk ploidy value that can be obtained from other NGS technologies, but whether the cell had two, four, or more than four sets of chromosomes. Thus, the development of a deep learning-based model able to quantify bulk ploidy complementary to NGS-derived ploidy can be important to further strengthen the predicted ploidy. In addition, it is known that ploidy patterns are cancer type-specific, and cancers that originate from related tissues tend to exhibit similar ploidy patterns. Thus, having a pan-cancer model might be valuable, allowing the model to learn common ploidy patterns across tissues^3,24^, increasing its potential for broader clinical use.

In this work, we present **Ploi**dy **Vi**sion **T**ransformer (PloiViT), a vision transformer-based model for the digital quantification of ploidy from pan-cancer H&E digital pathology slides. Furthermore, we compare the performance with a traditional ML model, XGBoost, and also with a late fusion model that fuses the outputs of both models. We make use of a WSI representation based on the aggregation of features obtained using two different feature extractors: an ImageNet pre-trained ResNet50, and CTransPath^19^ (defined as ResNet and CTrans going forward), given the satisfactory results that these models have showed in other tasks. We train and validate all models using fifteen different cancer types, and compare the predictions between them and between the feature extractors. Furthermore, we show that transformer-based models outperform traditional ML models when tested on independent cohorts. Finally, ploidy predictions are significantly different between important phenotypes (gleason score in prostate cancer, IDH-1-mutation in glioblastoma and HER2 positive in breat cancer).

## Results

### Machine learning models accurately predict ploidy values from digital pathology slides

We used TCGA as our discovery cohort to develop our models (see Methods). From this cohort, we first used an 80-20% training test split, and used the 80% training data for model development. To validate and select the different hyperparameters of the models, we performed a five-fold cross-validation over the training set. We compared the performance of three different models, using features from two different feature extractors: ResNet and CTrans. All details about model training can be found in the Methods section.

Firstly, we computed the performance across the five test sets in the five-fold cross-validation. For all performance metrics, root mean squared error (RMSE), mean absolute error (MAE), and mean absolute percentage error (MAPE), the model that combines the outputs from PloiViT and XGBoost (PloiViT+XGBoost) using CTrans features outperformed the rest of models in TCGA samples (Table 1). Models using CTrans features reached a superior performance in comparison to those using ResNet features. This is especially clear for the PloiViT model, where there is a difference of 0.2568 in terms of RMSE, and there is a significant difference between the model trained using ResNet features and that trained with CTrans features (paired t-test p-value ≤ 0.01). Furthermore, we computed all metrics per cancer type by joining the predictions over all test sets, simulating the performance over the whole training split, to observe the difference in performance between them (Figure 2 A, Supplementary Tables 1, 2, and 3). The best results were obtained for TCGA-PRAD (RMSE 0.3148), TCGA-THCA (RMSE 0.3346), and TCGA KIRP (RMSE 0.3346) in terms of RMSE. The worse results were obtained in TCGA-LUAD (RMSE 0.9867), TCGA-STAD (RMSE 0.9020), and TCGA-BRCA (RMSE 0.8164).

**Table 1.** Machine learning models accurately predict ploidy on TCGA. Metrics obtained per model, and feature extractor in the five-fold cross validation. The fusion model that uses CTrans features is the one that obtains the best results. Using CTrans features outperformed using ResNet features for both models. Best results are highlighted in bold.

| Model | RMSE (mean ± std) | MAE (mean ± std) | MAPE (mean ± std) |
| --- | --- | --- | --- |
| PloiViT ResNet | 1.0349 ± 0.1294 | 0.8540 ± 0.1202 | 0.3695 ± 0.0779 |
| XGBoost ResNet | 0.7496 ± 0.1071 | 0.6396 ± 0.0524 | 0.2563 ± 0.0159 |
| PloiViT+XGBoost ResNet | 0.8241 ± 0.1011 | 0.6881 ± 0.0800 | 0.2922 ± 0.0439 |
| PloiViT CTrans | 0.7781 ± 0.0997 | 0.6078 ± 0.0945 | 0.2439 ± 0.0362 |
| XGBoost CTrans | 0.7203 ± 0.1174 | 0.5929 ± 0.0702 | 0.2358 ± 0.0169 |
| PloiViT+XGBoost CTrans | **0.7002 ± 0.0903** | **0.5599 ± 0.0622** | **0.2241 ± 0.0172** |

Once we selected the different hyperparameters in the five-fold cross-validation (see Methods), we trained all models on the 80% training data, using 20% of it as validation for early stopping purposes. Then, we evaluated the model on the hold-out test set (Figure 2B, Table 2, Supplementary Tables 4, 5, and 6). Again, the combination of PloiViT and XGBoost using CTrans features outputs outperformed the rest of the models in terms of MAE and MAPE, reaching a RMSE of 0.7221, a MAE of 0.4730, and a MAPE of 0.1694. Only, XGBoost using CTrans features outperformed the fusion model in terms of RMSE (0.6810). This maintains the superiority of the CTrans over the ResNet features, and also the fusion of both models. A comparison between the real values and predictions per cancer type is presented in Supplementary Figure 1.

**Table 2.** Machine learning models generalize to the TCGA hold-out test set. Metrics obtained per model, and feature extractor in the TCGA hold-out test set. The fusion model that uses CTrans features is the one that obtains the best results. Using CTrans features outperformed using ResNet features for both models. Best results are highlighted in bold.

| Model | RMSE | MAE | MAPE |
| --- | --- | --- | --- |
| PloiViT ResNet | 0.8226 | 0.6710 | 0.3007 |
| XGBoost ResNet | 0.7007 | 0.5743 | 0.2400 |
| PloiViT+XGBoost ResNet | 0.7318 | 0.5972 | 0.2605 |
| PloiViT CTrans | 0.9110 | 0.6592 | 0.2353 |
| XGBoost CTrans | **0.6810** | 0.5249 | 0.2189 |
| PloiViT+XGBoost CTrans | 0.7221 | **0.4730** | **0.1694** |

### Transformer-based models demonstrate superior generalization in independent cohorts

We evaluated our models on two independent cohorts, CPTAC^25^ and PBTA^26^. For CPTAC, we assessed performance on LUAD and GBM using models trained on TCGA and the two feature extractors (ResNet and CTrans). In terms of MAE and MAPE, PloiViT with CTrans features consistently outperformed other models, yielding a CPTAC-GBM MAE of 0.5499 and a CPTAC-LUAD MAE of 0.9216 (Table 3, Supplementary Tables 7, 8). For RMSE, PloiViT CTrans was superior for CPTAC-LUAD, while the PloiViT+XGBoost fusion model with ResNet features narrowly edged out PloiViT CTrans for CPTAC-GBM (0.8354 vs. 0.8333). Notably, these results mirrored TCGA performance, where PloiViT CTrans achieved RMSEs of 0.9569 for TCGA-GBM and 1.5387 for TCGA-LUAD on the hold-out test set (Table 2).

**Table 3.** Root mean squared error (RMSE) obtained per model and per cancer type in the CPTAC cohort. Best results are highlighted in bold.

| CPTAC Project | PloiViT |  | XGBoost |  | PloiViT+XGBoost |  |
| --- | --- | --- | --- | --- | --- | --- |
|  | ResNet | CTrans | ResNet | CTrans | ResNet | CTrans |
| CPTAC-GBM | 0.8458 | 0.8354 | 0.8406 | 0.8525 | **0.8333** | 0.8378 |
| CPTAC-LUAD | 1.3090 | **1.1804** | 1.3215 | 1.2590 | 1.3128 | 1.2128 |

Subsequently, we predicted ploidy values for 125 PBTA slides (formed by four different cancer types) with determined ploidy (see Methods). PloiViT with CTrans features again excelled across all metrics, reaching an RMSE of 0.6758, MAE of 0.4792, and MAPE of 0.1649, surpassing XGBoost CTrans which obtained an RMSE of 0.7041, MAE of 0.5698, and MAPE of 0.2132 (Table 4). These low error values, especially the MAPE, underscore the strong generalization of our transformer-based model to unseen cohorts.

**Table 4.** Metrics obtained per model, and feature extractor in the PBTA dataset. Best results are highlighted in bold.

| Model | RMSE | MAE | MAPE |
| --- | --- | --- | --- |
| PloiViT ResNet | 0.7157 | 0.5523 | 0.1990 |
| XGBoost ResNet | 0.6915 | 0.6053 | 0.2358 |
| PloiViT+XGBoost ResNet | 0.6918 | 0.5773 | 0.2167 |
| PloiViT CTrans | **0.6758** | **0.4792** | **0.1649** |
| XGBoost CTrans | 0.7041 | 0.5698 | 0.2132 |
| PloiViT+XGBoost CTrans | 0.6811 | 0.5244 | 0.1890 |

### Prostate, glioblastoma, and breast cancer patients with specific phenotypes present higher predicted ploidy values

Next, we tested whether samples from patients with specific phenotypes presented higher predicted ploidy values than other patients. Specifically, we tested whether prostate cancer patients within a high risk group (based on the Gleason score) showed a higher ploidy value. Furthermore, we compared the predicted ploidy value between glioblastoma patients with and without the IDH-1 mutation. Finally, we tested the predicted ploidy value between breast cancer patients with negative and positive epidermal growth factor receptor (HER2). Thus, given the generalization performance of the PloiViT CTrans model, we used it to predict the ploidy values in all TCGA-PRAD (*N* = 895), TCGA-GBM (*N* = 236), and TCGA-BRCA (*N* = 1098) samples. It must be noted that the predictions were made for unseen samples, since only a portion of samples were present in the training set (*N* = 55 in the case if PRAD, *N* = 155 for GBM, and *N* = 90 for BRCA). Furthermore, during training our models were blind to any kind of phenotype stratification. For TCGA-PRAD samples, we divided the patients in three risk groups based on their Gleason Score (low risk: Gleason Group 1, intermediate risk: Gleason Groups 2, and 3, and high risk: Gleason Groups 4, and 5). For the TCGA-GBM samples, we split the patients in two groups: IDH-1 wildtype or mutant patients. Finally, for TCGA-BRCA we splitted the patients in two groups: HER2 negative and positive.

The high-risk patient group exhibited significantly higher predicted ploidy values compared to both low- and intermediate-risk groups (Mann-Whitney u test p-value ≤ 0.0001 in both cases, Figure 3 A). No significant difference was found between low- and intermediate-risk groups (Mann-Whitney u test p-value ≥ 0.05). Mean ploidy values (95% CI) were 2.1710 (2.1669-2.1920), 2.0666 (2.0722-2.1117), and 2.0703 (2.0774-2.0950) for high-, low-, and intermediate-risk groups, respectively.

Similarly, IDH-1 mutated patients showed significantly higher predicted ploidy values than wild-type patients (Mann-Whitney u test p-value ≤ 0.0001, Figure 3 B). Mean values (95% CI) were 2.4528 (2.3334-2.4631) and 2.2361 (2.2326-2.2827) for mutated and wild-type patients, respectively.

Additionally, HER2-positive patients exhibited significantly higher predicted ploidy values than HER2-negative patients (Mann-Whitney u test p-value ≤ 0.0001,, Figure 3 C), with mean values (95% CI) of 2.8770 (2.8506-2.9312) and 2.7516 (2.7600-2.7952), respectively.

### Transformer-based models can accurately predict tile-level spatial ploidy values across the tissue

We then used a spatial transcriptomics glioblastoma cohort^27^ (see Methods), to test whether our models could spatially predict ploidy values across the tissue at tile level. We used the pretrained models on TCGA, predicted the ploidy value for each spot, and compared with the NGS-derived ploidy value using the inferCNV R package. We computed the RMSE, MAE, and MAPE for each model across 16 slides, by using all spots predictions and NGS-derived values (Table 5, and Supplementary Tables 9, and 10).

**Table 5.** Root mean squared error (RMSE) obtained per model and slide in the spatial glioblastoma cohort. Given the low performance of the XGBoost model, we did not compute the performance of the PloiViT+XGBoost model. Best results are highlighted in bold.

| Slide ID | PloiViT |  | XGBoost |  |
| --- | --- | --- | --- | --- |
|  | ResNet | CTrans | ResNet | CTrans |
| 269_T | 0.1102 | **0.0540** | 0.6485 | 0.7929 |
| 270_T | 0.1451 | **0.0961** | 0.7061 | 0.7478 |
| 296_T | **0.1923** | 0.1973 | 0.5764 | 0.7851 |
| 275_T | 0.0704 | **0.0237** | 0.6886 | 0.5420 |
| 248_T | 0.0397 | **0.0154** | 0.6089 | 0.6642 |
| 334_T | 0.2668 | **0.0508** | 0.6592 | 0.6106 |
| 268_T | 0.1259 | **0.0633** | 0.6628 | 0.7511 |
| 255_T | **0.1563** | 0.1860 | 0.6154 | 0.6561 |
| 259_T | **0.0469** | 0.0602 | 0.6296 | 0.7280 |
| 266_T | **0.0367** | 0.1263 | 0.7280 | 0.7021 |
| 242_T | 0.1423 | **0.0736** | 0.6376 | 0.9478 |
| 262_T | 0.1264 | **0.1104** | 0.6947 | 0.6340 |
| 260_T | 0.0738 | **0.0272** | 0.5712 | 0.7858 |
| 243_T | **0.0355** | 0.0544 | 0.6046 | 0.7848 |
| 265_T | 0.2114 | **0.2101** | 0.6061 | 0.6265 |
| 304_T | **0.1896** | 0.1927 | 0.6328 | 0.6632 |
| Mean | 0.1231 | **0.0963** | 0.6419 | 0.7139 |
| STD | 0.0675 | **0.0645** | 0.0442 | 0.0940 |

PloiViT significantly outperformed the XGBoost model (paired t-test p-value ≤ 0.0001), both using the ResNet and the CTrans features (Table 5). Furthermore, PloiViT using CTrans features obtained the best mean performance (mean RMSE: 0.0963 ± 0.0645, mean MAE: 0.0859 ± 0.0597, and mean MAPE: 0.0429 ± 0.0298) closely followed by PloiVIT using ResNet features (mean RMSE: 0.1231 ± 0.0674, mean MAE: 0.1095 ± 0.0631, and mean MAPE: 0.0547 ± 0.0315). However, this difference was not significant between the two PloiViT models (paired t-test p-value ≥ 0.05). Error metrics obtained by XGBoost were worse, reaching a mean RMSE of 0.6419 and 0.7139 using ResNet and CTrans features respectively. This validates the previously obtained results in independent cohorts, showing that PloiViT presents better generalization capabilities to unseen data. In addition, the results obtained by PloiViT using CTrans features in terms of MAPE are comparable to those obtained in the hold-out test set by the PloiViT+XGBoost model using CTrans features (0.1143, Supplementary Table 10) in the TCGA-GBM cohort, further validating these results.

## Discussion

The study of DNA ploidy, which is highly correlated with aneuploidy, is a rapidly emerging focus in cancer research due to the aneuploidy links with prognosis and drug resistance. It promotes evolutionary flexibility, cellular adaptation, cancer stem-cell properties, and immune evasion, all key drivers of aggressive tumor behavior. Targeting these aneuploidy-linked mechanisms holds therapeutic promise, with early successes in exploiting synthetic lethality, mitotic defects, and aneuploidy-tolerance mechanisms^2,4^. Thus, having a reliable and cheap ploidy quantification tool that complements NGS-derived ploidy can advance patient treatment and clinical decisions.

In this work we have presented PloiViT, a transformer-based model capable of accurately quantifying ploidy from routinely obtained H&E stained digital pathology slides. We compared the performance of our model with XGBoost and a late fusion model. While within the TCGA cohort a better performance was obtained by the late fusion and ML models (Tables 1, 2), this performance was not maintained when tested on independent cohorts. PloiViT using CTrans features outperformed both XGBoost and the late fusion model on CPTAC in terms of MAE and MAPE (Supplementary Tables 7, 8), for all metrics in the case of the PBTA cohort (Table 4), and for all slides in the spatial GBM cohort (Table 5, and Supplementary Tables 9, and 10). Furthermore, it must be noted that PBTA contains cancer types outside those in the training set, showing impressive results.

These results show the better generalization performance that deep learning models have over traditional ML methods. We also hypothesize this could be due to the attention mechanism used in PloiViT. To train XGBoost, a single feature vector is used, so the average of all clusters centroids is used as input to the model. However, for PloiViT, all 100 centroid feature vectors are used as input, allowing the model to learn relationships between them thanks to the self-attention operation. Nevertheless, it is still surprising the high-accuracy that a traditional ML can achieve on this task, especially comparing the computational requirements to train both models.

If we compare the performance between the two feature extractors, it is clear that using a digital pathology specific model like CTrans provides an advantage over ResNet features, especially observing the results obtained both in the five-fold cross validation and in the hold-out test set (Tables 1, 2). This result has recently been reported in other works, and it seems to be the future path for digital pathology classification^28–30^. Testing the CTrans feature extractor in non-TCGA cohorts is crucial, since CTrans was trained using TCGA samples. When we tested the pre-trained models in the independent cohorts, the models using CTrans features also outperformed the ResNet models. This was not the case for all cancer types (e.g. TCGA-LUAD, TCGA-STAD, or TCGA-GBM, Supplementary Table 4), but there was an improvement in the mean performance of the models. Thus, similarly to what other researchers have reported in relevant cancer tasks, using a SSL trained model on digital pathology as a feature extractor outperforms feature extractors trained on natural images for ploidy quantification^20,21^.

To further study whether our predictions accurately represented the effect of some phenotypes in the samples, we compared the predicted ploidy value in three TCGA cohorts (PRAD,GBM, and BRAC), for the gleason risk group, the IDH-1 mutation respectively, and the HER2 status respectively. It must be noted that we predicted the value for all available samples, not restricting ourselves to the handful of samples for which the NGS-derived ploidy value is known. For TCGA-PRAD, we showed that a significantly higher ploidy value was predicted within the high risk group in comparison to the low and intermediate risk groups (Figure 3 A). The gleason score inform us about the prognostic perspective of the patient and also helps guide the patient’s therapy^31–33^. High risk patients group are those with advanced neoplasms that are unlikely to be cured. Given the association between abnormal ploidy values and prognosis^34^, it is expected that PRAD patients within the high risk group present a higher mean ploidy value than those in early stages of the disease. Next, we also compared the predicted ploidy value in TCGA-GBM samples between wildtype and those samples with a mutation in IDH-1. Mutations affecting the DNA damage response (e.g. IDH-1 mutation in GBM) enable polyploid cells to further proliferate^35^. Thus, a higher predicted ploidy value is expected in those samples with the IDH-1 mutation, which was the case in our experiments (Figure 3 B). Finally, HER2 overexpression or amplification in BRCA is a marker of aggressive disease and poorer outcomes. This receptor activates signaling pathways that drive cancer cell growth and proliferation^36^. Thus, given the higher proportion of cancer cells, it is expected that HER2 positive patients have a higher ploidy value than negative patients. This was the case in our experiments, where a significant difference was found in the predicted ploidy between positive and negative patients (Figure 3 C). These experiments validate the capability of our model to predict ploidy values that accurately align to the expected biology of important phenotypes and disease stages.

Finally, even though our model was trained for slide-level prediction, we decided to test the model capabilities by predicting spatial patterns of ploidy values across the tissue. By using a spatial GBM cohort, we showed that PloiViT models outperform XGBoost by a large margin, and using the CTrans features also outperform ResNet features (Table 5). Furthermore, it is noticeable that the error obtained by PloiViT is low, and comparable to that obtained by the late fusion model in the TCGA-GBM hold-out test set cohort (Supplementary Table 4). It must be noted that the tiles required upsampling to the desired resolution, since the spatial spot comprehends a tile of approximately 56 × 56 pixels. Furthermore, XGBoost allows to use a single tile to make the prediction, while for PloiViT we are using a sliding-window method (see Methods), which might affect the performance of one over the other. Nevertheless, PloiViT outperformed XGBoost in the rest of independent cohorts, therefore, it is expected that the same happens in the spatial cohort.

Our models demonstrate the ability of deep learning to quantify ploidy across multiple cohorts. However, there are important considerations to be made. Our models were trained exclusively on TCGA data, and while this is an invaluable resource, incorporating data from multiple sources would enhance model diversity. Additionally, since our models’ quantification is trained on the TCGA wider-scale validation using different quantification methods is necessary. Furthermore, we currently quantify ploidy at the bulk level due to data limitations. A high-quality, H&E-matched spatial transcriptomics dataset at single-cell resolution would be transformative, enabling the development of a model capable of predicting ploidy for individual cells. While such datasets are not yet available, they hold the potential for significant advancements in this field.

PloiViT demonstrates the superiority of transformer-based models over traditional machine learning approaches for ploidy prediction, offering improved generalization and accuracy. As a crucial addition to the digital quantification toolkit, PloiViT has the potential to derive critical clinical information from cost-effective modalities, positively impacting patient care. We foresee that PloiViT can be used to complement NGS-derived ploidy, providing a more robust final quantification by relaying on both methodologies. To facilitate the use of the tool and further research, we have made our models available both in the source code, a user-interface inference application, and a python package (see Code Availability). Future research directions include extending our model to additional cancer types, subject to data availability. Integrating multiple datasets, given sufficient samples, would further enhance performance. Moreover, the availability of spatial transcriptomics data would enable training a model with improved accuracy and the ability to pinpoint spatial patterns of ploidy.

## Methods

### Patient cohorts

#### The Cancer Genome Atlas Project

For model training and evaluation, anonymized patient data were retrieved from the publicly available The Cancer Genome Atlas (TCGA) archive (available at https://portal.gdc.cancer.gov). We used paraffin-embedded (FFPE) whole slide images (WSIs) and ploidy values from fifteen cancer types, including bladder urothelial carcinoma (BLCA), breast invasive carcinoma (BRCA), blioblastoma multiforme (GBM), head and neck squamous cell carcinoma (HNSC), kidney chromophobe (KICH), kidney renal clear cell carcinoma (KIRC), kidney renal papillary cell carcinoma (KIRP), liver hepatocellular carcinoma (LIHC), lung adenocarcinoma (LUAD), lung squamous cell carcinoma (LUSC), prostate adenocarcinoma (PRAD), skin cutaneous melanoma (SKCM), stomach adenocarcinoma (STAD), Thyroid carcinoma (THCA), and uterine corpus endometrial carcinoma (UCEC). The number of patients and WSIs is listed in Supplementary Table 11.

#### Clinical Proteomic Tumor Analysis Consortium

For validation, anonymized patient data were retrieved from the publicly available Clinical Proteomic Tumor Analysis Consortium (CPTAC) cohort^25^ (https://www.cancerimagingarchive.net/collections). We downloaded matched WSIs from two cancer types, lung adenocarcinoma (LUAD) and glioblastoma multiforme (GBM). The sample size is described in Supplementary Table 12.

#### Pediatric Brain Tumor Atlas

Anonymized WSIs from the pediatric brain cancer cohort from PBTA consisted of four tumor subtypes: low-grade glioma/astrocytoma (LGG), high-grade glioma/astrocytoma (HGG), Ganglioglioma, others and others (https://kidsfirstdrc.org). Clinical data was downloaded from the original publication of Shapiro et al.^26^. Ploidy values were inferred via Control-FREEC^37^. The sample size is described in Supplementary Table 13.

#### Spatial Glioblastoma Dataset

Spatial transcriptomic data and matched histology images of GBM were obtained from a published study by Ravi et al.^27^ (https://datadryad.org/stash/dataset/doi:10.5061/dryad.h70rxwdmj). Ploidy values per spot were obtained using the inferCNV package^38^.

### Preprocessing of Whole Slide Images

Whole-slide images (WSIs) were acquired in SVS format and downsampled to a magnification of 20x (0.5 *µ*m per pixel). The Otsu threshold method was applied to generate a tissue mask, facilitating the exclusion of tiles predominantly containing white background^39^. Each WSI was partitioned into non-overlapping tiles measuring 256×256 pixels (128 *µ*m x 128 *µ*m). For each slide, a maximum of *N* = 4000 tiles was randomly selected, filtering out those containing more than 20% background and tiles with low contrast.

For slide-level feature representation, the selected tiles were grouped into bags. Initially, a ResNet-50 module pretrained on ImageNet, or the pre-trained CTransPath model, were employed to transform each tile into a feature vector (1*xN*, being *N* the embedding dimension of each feature extractor, *N* = 2048, 768). Subsequently, the k-means algorithm was utilized to cluster similar tiles within each slide into *K* = 100 clusters. The features of patches within each cluster were then averaged, yielding a matrix of 100*xN* or vectors representing the slide. This allows to capture the intrinsic characteristics of groups of tiles while reducing the computational requirements for a transformer-based model.

### Machine learning models details

In this work we have used two different models: PloiViT and XGBoost. PloiViT draws inspiration from the original ViT architecture proposed by Dosovitskiy et al.^22^, which extends the Transformer framework from natural language processing (NLP) to the realm of computer vision^40^. Furthermore, it is also based on *SEQUOIA*, our proposal for digital quantification of cancer transcriptomes^41^. In the ViT framework, images are segmented into patches of size 16 × 16 pixels, akin to “tokens” in NLP. These patches undergo linear projection to extract feature vectors, which are subsequently processed by a transformer encoder. This encoder refines the input representation and directs it to a multi-layer perceptron (MLP) head for final prediction. A schematic representation of the architecture can be observed in Figure 1 A.

**Figure 1.**
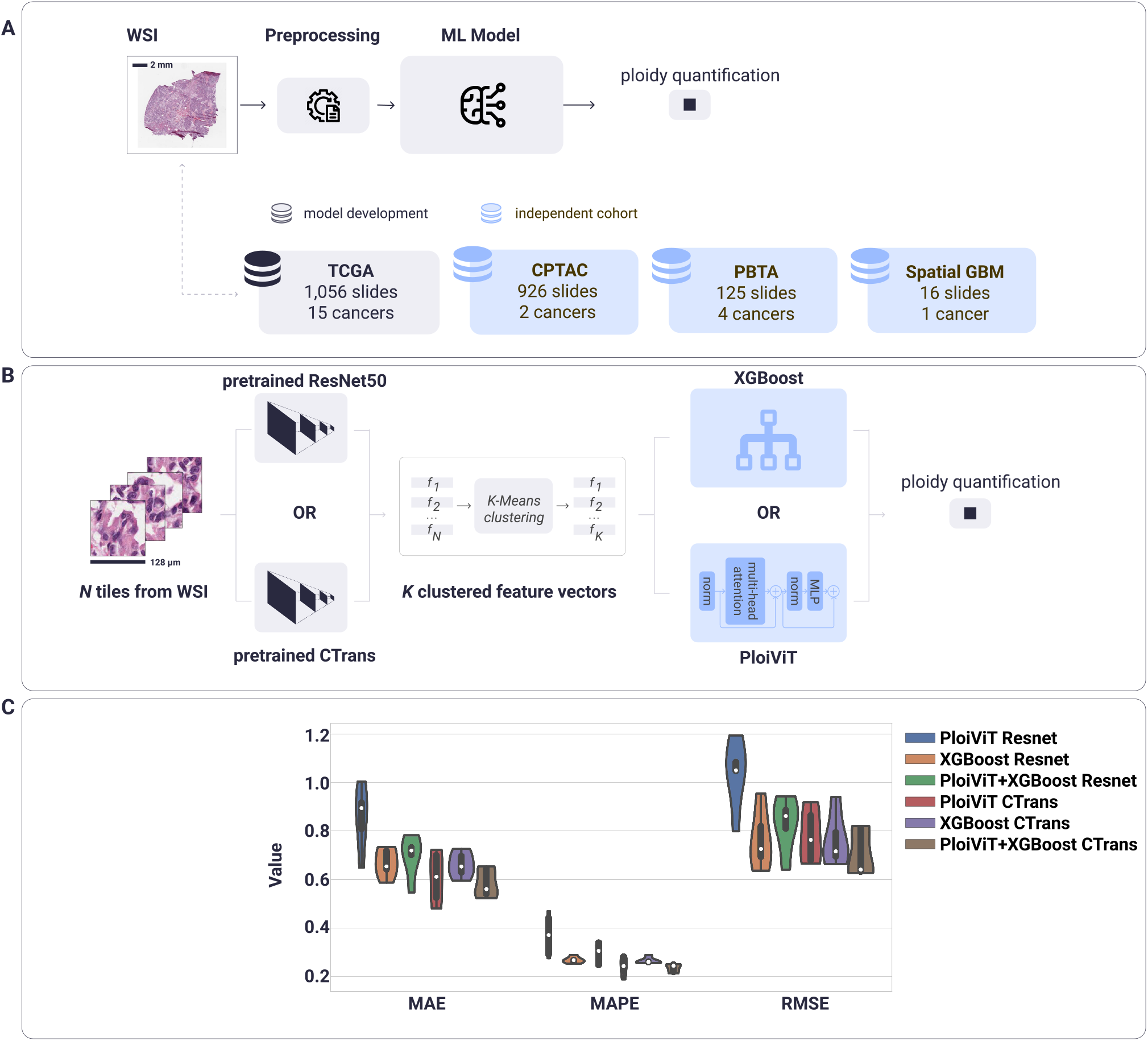
Machine learning for accurate ploidy quantification from digital pathology slides. **Panel A**: Our models use as input a whole-slide image, applies several preprocessing steps (see Methods) and predict the overall ploidy value. We used TCGA samples from fifteen cancer types as our discovery set, and tested our model on three independent cohorts for survival prediction and spatial validation. **Panel B:** Oue training and prediction pipeline. It uses as input N tiles from the whole-slide image, and a pretrained Resnet50 or CTrans model is used to obtain N feature vectors. A K-Means clustering algorithm is used to reduce the N features vectors to K clusters, and the centroid vector per cluster is computed. These K centroid feature vectors are used as input to the transformer encoder or the XGBoost model to perform the ploidy prediction. **Panel C:** We compared the performance of PloiViT, XGBoost, and the fusion of both models (named PloiViT+XGBoost) using either ResNet50 or CTrans features in a five-fold cross validation. In this plot, we represent the performance of each model in terms of mean absolute error (MAE), root mean squeared error (RMSE), and mean absolute percentage error (MAPE). The lower the values, the better the performance. The best median performance was achieved by PloiViT+XGBoost using CTrans features, followed by XGBoost using CTrans features.

**Figure 2.**
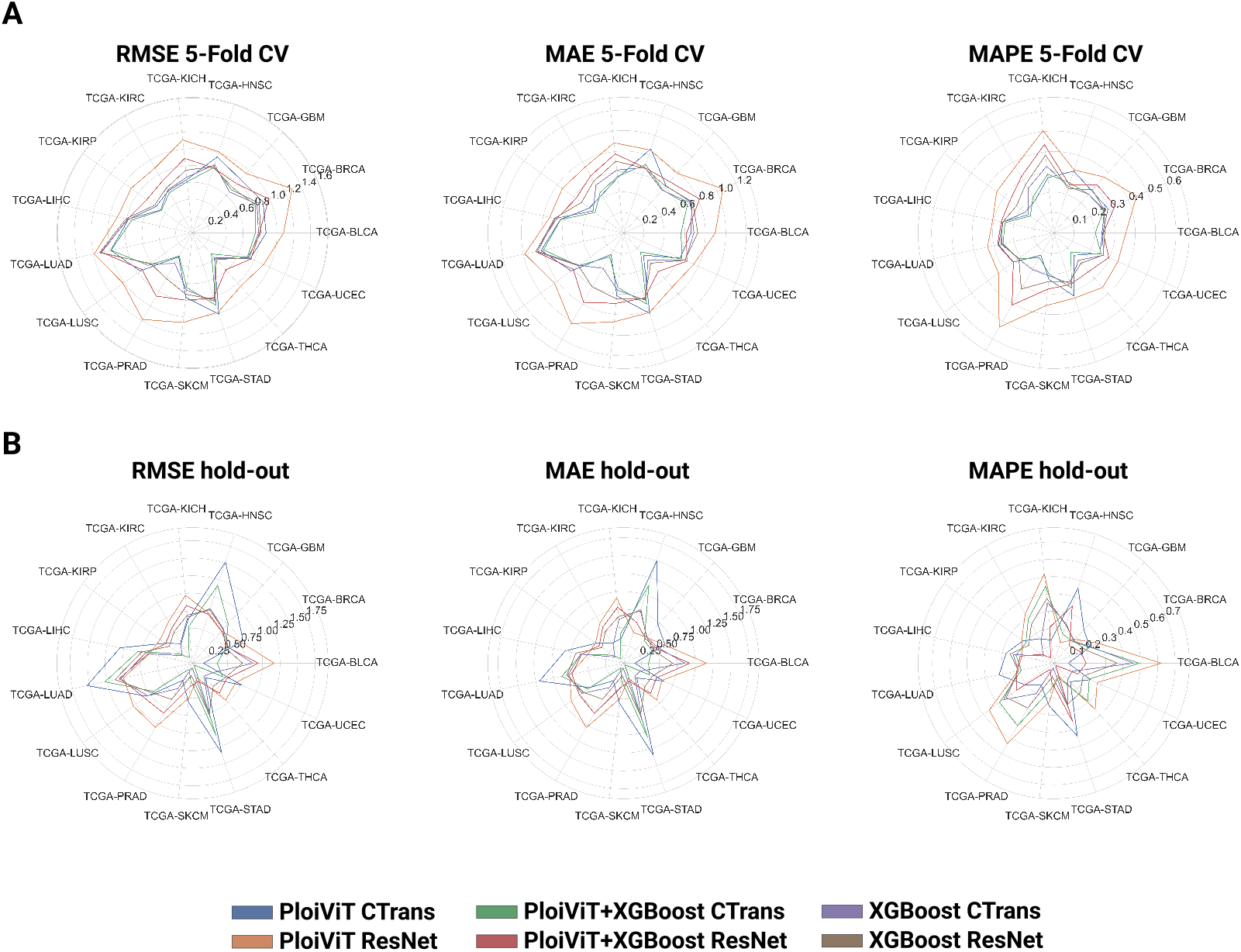
Models using CTrans features outperform models using ResNet features for pan-cancer ploidy prediction. **Panel A:** Comparison in terms of root mean squared error (RMSE), mean absolute error (MAE), and mean absolute percentage error (MAPE) between PloiViT, XGBoost, and PloiViT+XGBoost models using ResNet and CTrans features. For the metrics, the closer to zero denotes a better performance. These results were obtained by joining the predictions in all test sets in the five-fold cross validation of the discovery set (TCGA). **Panel B:** Comparison in terms of RMSE, MAE, and MAPE between PloiViT, XGBoost, and PloiViT+XGBoost models using ResNet and CTrans features in the TCGA hold-out test set.

**Figure 3.**
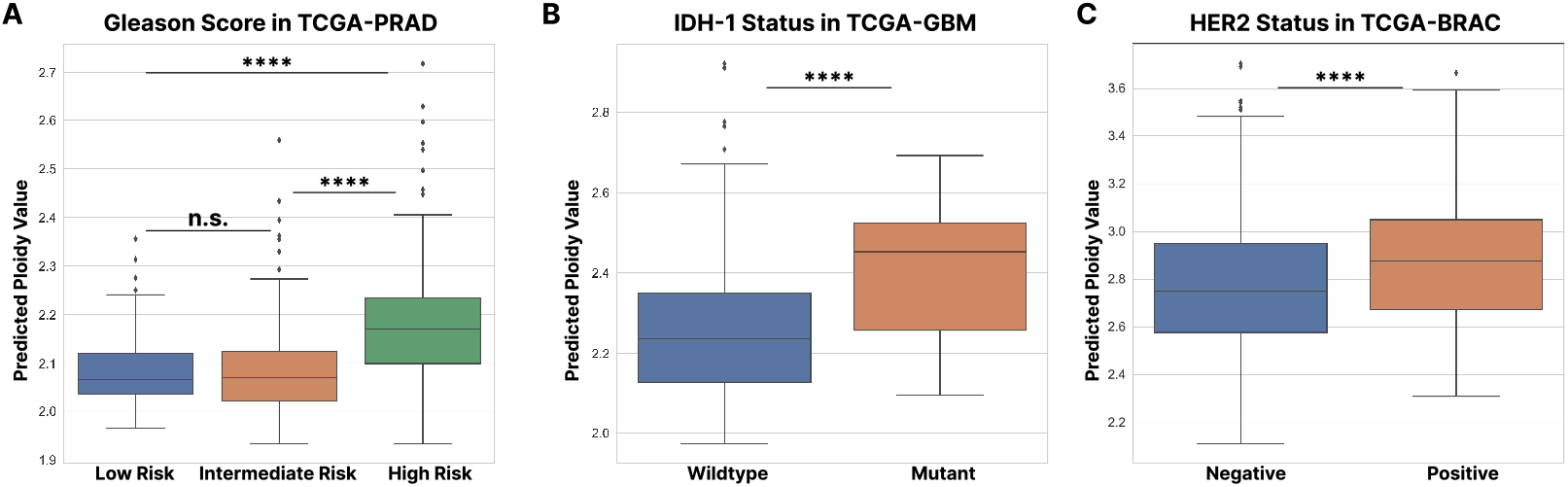
Prostate cancer within a higher gleason risk group and glioblastoma patients with IDH-1 mutation present a higher predicted ploidy. **Panel A:** Difference in predicted ploidy value between risk groups in TCGA-PRAD defined by their gleason score. Patients in the high risk group showed a significantly different predicted ploidy value with low and intermediate risk patients. **Panel B:** TCGA-GBM IDH-1 mutant patients presented a significantly higher predicted ploidy value than wildtype patients. **Panel C:** TCGA-BRCA HER2 positive patients presented a significantly higher predicted ploidy value than HER2 negative patients. n.s. denotes no significance (p-value ≥ 0.05), **** (p-value ≤ 0.0001).

In our approach, the image segmentation concept aligns with the partitioning of WSIs into *K* feature clusters, with a feature extractor dimensionality of *N* (*K* = 100, *N* = 2048, 768). The resulting 100 × *N* feature matrix is then channeled into the transformer encoder, comprising 6 encoder blocks, 16 attention heads, and a head dimension of 64. This transformer encoder facilitates the exploration of inter-cluster relationships, crucial for determining their relevance in slide-level prediction tasks. Subsequent to layer normalization, the transformed output is conveyed to an MLP layer with dimensions *N* × 1, to predict the ploidy value.

For XGBoost, we used the implementation provided in the XGBoost Python Package. We used a Grid Search Cross-Validation to find the best hyperparameters. We defined the following set of hyperparameters for the search: *min_child_weight*: [1, 5, 10], *gamma*: [0.5, 1, 1.5, 2, 5], *subsample*: [0.6, 0.8, 1.0], *colsample_bytree*: [0.6, 0.8, 1.0], *max_depth*: [3, 4, 5, 6], and *eta*: [0.1, 0.3, 0.6].

### Training and evaluation details

For training and evaluation of the model, we conducted a five-fold cross-validation using data from the TCGA cohort. Firstly, the data was partitioned at patient-level in a 80%-20% training and hold-out test sets. The hold-out test set was not used during model training, and it was only used for evaluating the model after the hyperparameter selection and model training. Then, a five-fold cross-validation was used within the training set. In each fold *i*, the dataset was partitioned on patient level, allocating 80% for ‘global’ training and 20% for testing. To determine the optimal stop point for training the model in fold *i*, the ‘global’ training set *i* was further split into 90% for training and 10% for internal validation. We used the Mean Squared Error (MSE) as the loss function during model training, with each model being trained for a maximum of 100 epochs. For early stopping and determining the point for model saving, we used the MSE on the validation set, which a patience value of 20 epochs. Throughout this process, we used a fixed learning rate of 3 × 10^−3^ and batch size of 32, and the model parameters were optimized with the Adam optimizer. Then, once the best hyperparameters were selected, the whole training set (80% of the data) was used to train both models model using the same hyperparameters and optimizer in the case of PloiViT, and then the predictive performance was tested on the hold-out test set. For validating the models, we computed the MSE values, the Mean Average Error (MAE) and the Mean Absolute Percentage Error (MAPE).

### Spatial prediction of ploidy value on tile level

To predict ploidy values at the tile level, we used two different approaches. For XGBoost we used the features from the tile to make the prediction. On the other hand, for *PloiViT*, we implemented a sliding-window technique. Beginning from the upper-left corner of the whole-slide image (WSI), we consider a window of size 10 × 10 tiles. To specify the window’s position, we denote it by the (*x, y*) coordinates of its upper-left tile. Consequently, the window in the upper-left corner is denoted as *w*_0,0_ (with the *x, y* axes following image processing conventions, where the origin is in the upper-left corner, and the x-axis increases to the right while the y-axis increases downwards).

The feature vectors of the tiles within window *w*_*x,y*_, which form a 100 × 2048 matrix, are fed into the model. Each individual tile’s feature vector serves as a ‘cluster mean vector’. The resultant predicted ploidy value 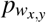 is recorded for all tiles in the window. Subsequently, the window is shifted to the right by a distance of *stride* tiles (*w*_*x*+*stride,y*_), and the predicted ploidy for each tile in this new window is saved. Once the window reaches the end of a row (i.e., when *x* + *stride* + 10 equals the width of the image), a new window begins *stride* tiles below the previous row (*w*_0,*y*+*stride*_). After traversing the entire WSI, the prediction for each tile is computed as the average of all values saved for that tile when it was part of a window *w*_*x,y*_. In our implementation, we opted for *stride* = 1 (larger strides entail less computational time but offer reduced granularity).

### Software details

All code was developed using Python 3.11. Models were trained on a single NVIDIA V100 with 80 GB RAM. As the deep learning framework we used pytorch^42^, and for computing the evaluation metrics, we used the scikit-learn python package^43^. Data visualization and data I/O was performed using the matplotlib, seaborn and pandas python libraries. *PloiViT* is encapsulated as a python package (*ploidyml*), that can be installed using pip, as described in the Code Availability section. Model weights are provided in the Github respository. The user-interface software was developed using the Streamlit python library. All requirements and library versions can be found in the *requirements*.*txt* file provided in the Github repository of the project.

## Acknowledgements

The results published here are in whole or in part based on data generated by the TCGA Research Network (https://www.cancer.gov/tcga).

M.P. was supported by the Belgian American Educational Foundation and FWO (grant number 1161223N). The research reported here was further supported by the National Cancer Institute (NCI) under award: R01 CA260271. The content is solely the authors’ responsibility and does not necessarily represent the official views of the National Institutes of Health.

## Author contributions statement

F.C.-P., N.A. and O.G. conceived and designed the study. F.C.-P., and E.M.C. performed data preprocessing. F.C.-P. and M.P. developed the code. N.A. and E.M.C. performed the analysis of the clinical impact. E.M.C. obtained the spatial ploidy values. F.C.-P. and M.P. generated the figures. O.G. and N.A. supervised the work and obtained the funding. F.C.-P. and O.G. wrote the manuscript with contributions and/or revisions from all authors. All authors reviewed the manuscript.

## Code Availability

The code used in this work and how to install the user-interface can be found in the following Github repository: https://github.com/gevaertlab/Ploidy-ML.

## Data Availability

Anonymized WSIs, and clinical data of the The Cancer Genome Atlas’ (TCGA) cohorts were retrieved from the publicly available Genomic Data Commons (GDC) portal (https://portal.gdc.cancer.gov). Ploidy values for model’s training were obtained from Supplementary Material Table 3 of the The ICGC/TCGA Pan-Cancer Analysis of Whole Genomes Consortium publication^44^. Anonymized WSIs from the Clinical Proteomic Tumor Analysis Consortium (CPTAC) cohort were were obtained from the Cancer Image Archive with the accession URL (https://www.cancerimagingarchive.net/collections). Survival data for TCGA and CPYAC and ploidy values for the CPTAC datasets were obtained from the the LinkedOmics platform (https://www.linkedomics.org/login.php). The pediatric brain tumor cohort from the Pediatric Brain Tumor Atlas (PBTA) available through the Gabriella Miller Kids First Data Resource Portal (https://kidsfirstdrc.org). Spatial transcriptomic data and matched histology images of GBM were obtained from a published study by Ravi et al. (https://datadryad.org/stash/dataset/doi:10.5061/dryad.h70rxwdmj).

## Additional information

To include, in this order: **Accession codes** (where applicable); **Competing interests** (mandatory statement).

The corresponding author is responsible for submitting a [competing interests statement](http://www.nature.com/srep/policies/index.html#competing) on behalf of all authors of the paper. This statement must be included in the submitted article file.

## References

1. Taylor, A. M. et al. Genomic and functional approaches to understanding cancer aneuploidy. Cancer cell 33, 676–689 (2018).

2. Simonetti, G., Bruno, S., Padella, A., Tenti, E. & Martinelli, G. Aneuploidy: Cancer strength or vulnerability? Int. journal cancer 144, 8–25 (2019).

3. Ben-David, U. & Amon, A. Context is everything: aneuploidy in cancer. Nat. Rev. Genet. 21, 44–62 (2020).

4. Vasudevan, A. et al. Single-chromosomal gains can function as metastasis suppressors and promoters in colon cancer. Dev. Cell 52, 413–428 (2020).

5. Leung, W. & Lao, T. Rapid aneuploidy testing, traditional karyotyping, or both? The Lancet 366, 97–98 (2005).

6. Augusto Corrêa dos Santos, R., Goldman, G. H. & Riaño-Pachón, D. M. ploidyngs: visually exploring ploidy with next generation sequencing data. Bioinformatics 33, 2575–2576 (2017).

7. Merkel, D. E. & McGuire, W. L. Ploidy, proliferative activity and prognosis. dna flow cytometry of solid tumors. Cancer 65, 1194–1205 (1990).

8. Silvestrini, R. et al. Flow cytometric analysis of ploidy in colorectal cancer: a multicentric experience. Br. journal cancer 67, 1042–1046 (1993).

9. Truong, K. et al. Fluorescence-based analysis of dna ploidy and cell proliferation within fine-needle samplings of breast tumors: A new approach using automated image cytometry. Cancer Cytopathol. Interdiscip. Int. J. Am. Cancer Soc. 84, 309–316 (1998).

10. Faggioli, F., Vezzoni, P. & Montagna, C. Single-cell analysis of ploidy and centrosomes underscores the peculiarity of normal hepatocytes. PloS one 6, e26080 (2011).

11. Cornish, T. C., Swapp, R. E. & Kaplan, K. J. Whole-slide imaging: routine pathologic diagnosis. Adv. anatomic pathology 19, 152–159 (2012).

12. Pantanowitz, L. et al. Review of the current state of whole slide imaging in pathology. J. pathology informatics 2, 36 (2011).

13. Madabhushi, A. & Lee, G. Image analysis and machine learning in digital pathology: Challenges and opportunities. Med. image analysis 33, 170–175 (2016).

14. Lu, M. Y. et al. Ai-based pathology predicts origins for cancers of unknown primary. Nature 594, 106–110 (2021).

15. Steyaert, S. et al. Multimodal data fusion for cancer biomarker discovery with deep learning. Nat. Mach. Intell. 5, 351–362 (2023).

16. Pizurica, M. et al. Whole slide imaging-based prediction of tp53 mutations identifies an aggressive disease phenotype in prostate cancer. Cancer Res. CAN–22 (2023).

17. Lu, M. Y. et al. Data-efficient and weakly supervised computational pathology on whole-slide images. Nat. biomedical engineering 5, 555–570 (2021).

18. Campanella, G. et al. Clinical-grade computational pathology using weakly supervised deep learning on whole slide images. Nat. medicine 25, 1301–1309 (2019).

19. Wang, X. et al. Transformer-based unsupervised contrastive learning for histopathological image classification. Med. image analysis 81, 102559 (2022).

20. Filiot, A. et al. Scaling self-supervised learning for histopathology with masked image modeling. medRxiv 2023–07 (2023).

21. Chen, R. J. et al. Towards a general-purpose foundation model for computational pathology. Nat. Medicine 1–13 (2024).

22. Dosovitskiy, A. et al. An image is worth 16×16 words: Transformers for image recognition at scale. arXiv preprint arXiv:2010.11929 (2020).

23. Wen, Z. et al. Deep-learning-based hepatic ploidy quantification using h&e histopathology images. Genes 14, 921 (2023).

24. Shukla, A. et al. Chromosome arm aneuploidies shape tumour evolution and drug response. Nat. communications 11, 449 (2020).

25. Edwards, N. J. et al. The cptac data portal: a resource for cancer proteomics research. J. proteome research 14, 2707–2713 (2015).

26. Shapiro, J. A. et al. Openpbta: The open pediatric brain tumor atlas. Cell Genomics (2023).

27. Ravi, V. M. et al. Spatially resolved multi-omics deciphers bidirectional tumor-host interdependence in glioblastoma. Cancer cell 40, 639–655 (2022).

28. El Nahhas, O. S. et al. Regression-based deep-learning predicts molecular biomarkers from pathology slides. Nat. communications 15, 1253 (2024).

29. Laleh, N. G. et al. Benchmarking weakly-supervised deep learning pipelines for whole slide classification in computational pathology. Med. image analysis 79, 102474 (2022).

30. Niehues, J. M. et al. Generalizable biomarker prediction from cancer pathology slides with self-supervised deep learning: A retrospective multi-centric study. Cell reports Medicine 4 (2023).

31. Egevad, L., Granfors, T., Karlberg, L., Bergh, A. & Stattin, P. Prognostic value of the gleason score in prostate cancer. BJU international 89, 538–542 (2002).

32. Helpap, B. et al. The significance of accurate determination of gleason score for therapeutic options and prognosis of prostate cancer. Pathol. & Oncol. Res. 22, 349–356 (2016).

33. Coward, J. & Harding, A. Size does matter: why polyploid tumor cells are critical drug targets in the war on cancer. Front. oncology 4, 123 (2014).

34. Bielski, C. M. et al. Genome doubling shapes the evolution and prognosis of advanced cancers. Nat. genetics 50, 1189–1195 (2018).

35. Shi, D. D., Anand, S., Abdullah, K. G. & McBrayer, S. K. Dna damage in idh-mutant gliomas: mechanisms and clinical implications. J. neuro-oncology 162, 515–523 (2023).

36. Loibl, S. & Gianni, L. Her2-positive breast cancer. The Lancet 389, 2415–2429 (2017).

37. Boeva, V. et al. Control-freec: a tool for assessing copy number and allelic content using next-generation sequencing data. Bioinformatics 28, 423–425 (2012).

38. Tickle, T., Georgescu, C., Brown, M. & Haas, B. Infer copy number variation from single-cell rna-seq data (2019).

39. Otsu, N. A threshold selection method from gray-level histograms. IEEE transactions on systems, man, cybernetics 9, 62–66 (1979).

40. Vaswani, A. et al. Attention is all you need. Adv. neural information processing systems 30 (2017).

41. Zheng, Y. et al. Digital profiling of cancer transcriptomes from histology images with grouped vision attention. BioRxiv (2023).

42. Paszke, A. et al. Pytorch: An imperative style, high-performance deep learning library. Adv. neural information processing systems 32 (2019).

43. Pedregosa, F. et al. Scikit-learn: Machine learning in python. J. machine Learn. research 12, 2825–2830 (2011).

44. of Whole Genomes Consortium, T. I. P.-C. A. Pan-cancer analysis of whole genomes. Nature 578, 82–93 (2020).

