## Supplementary Tables and Figure for "Towards Digital Quantification of Ploidy from Pan-Cancer Digital Pathology Slides using Deep Learning"

### ABSTRACT

### Supplementary Information

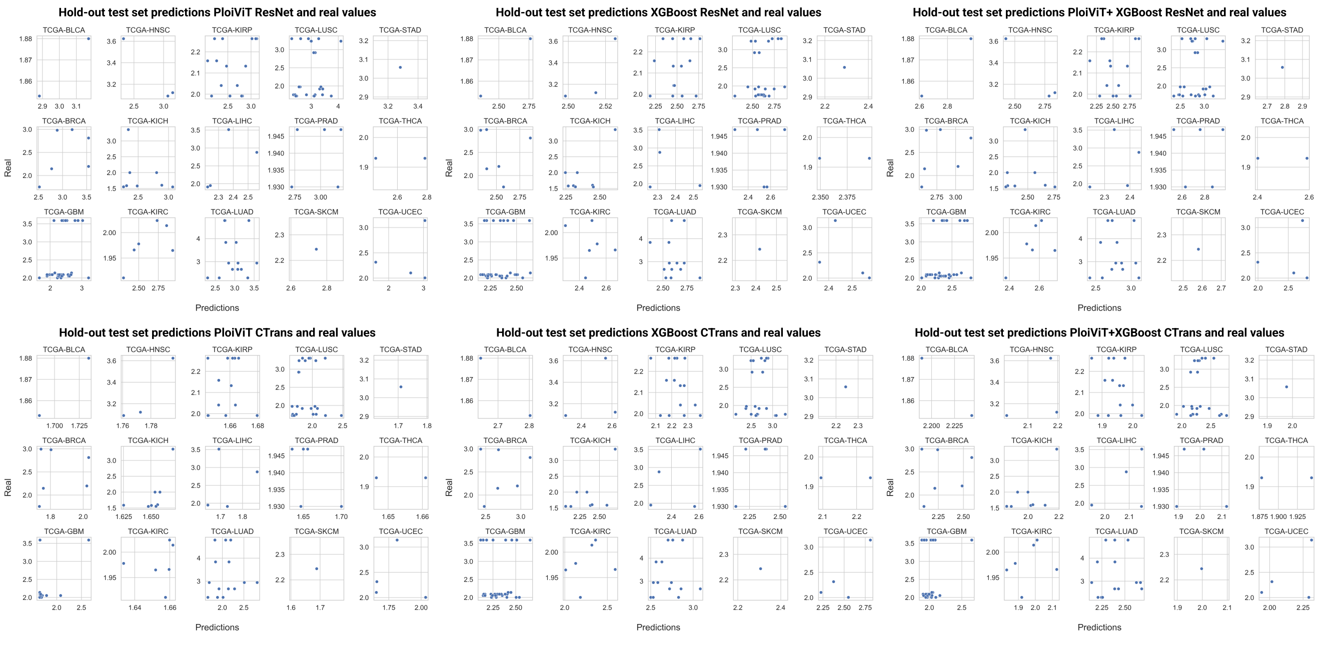


**Supplementary Figure 1.** Comparison between real values and predictions obtained per model in the TCGA hold-out test set.

**Supplementary Table 1.** Root mean squared error (RMSE) obtained per model and per cancer type in the five-fold cross-validation joining all test sets on TCGA samples. Best results are highlighted in bold.

| **TCGA Project** | **PloiViT** | | **XGBoost** | | **PloiViT+XGBoost** | |
|---|---|---|---|---|---|---|
| | ResNet | CTrans | ResNet | CTrans | ResNet | CTrans |
| TCGA-BLCA | 1.0795 | 0.8724 | 0.8076 | 0.7821 | 0.7932 | **0.7454** |
| TCGA-BRCA | 1.2910 | 0.9139 | 0.8652 | 0.8375 | 0.9909 | **0.8164** |
| TCGA-GBM | 0.9493 | 0.8525 | 0.6893 | **0.6552** | 0.7829 | 0.7166 |
| TCGA-HNSC | 1.0116 | 0.9475 | 0.8285 | 0.8509 | **0.8099** | 0.8141 |
| TCGA-KICH | 1.1087 | 0.6356 | 0.7479 | 0.6636 | 0.8891 | **0.6104** |
| TCGA-KIRC | 0.8907 | 0.5758 | 0.6170 | 0.5873 | 0.7116 | **0.5502** |
| TCGA-KIRP | 0.8999 | 0.5332 | 0.5577 | 0.4659 | 0.67157 | **0.45524** |
| TCGA-LIHC | 0.9085 | 0.7764 | 0.8081 | 0.7822 | 0.7887 | **0.7343** |
| TCGA-LUAD | 1.1901 | 0.9915 | 1.1258 | 1.1134 | 1.1020 | **0.9867** |
| TCGA-LUSC | 1.0165 | 0.7670 | **0.6159** | 0.6219 | 0.73916 | 0.6524 |
| TCGA-PRAD | 1.1847 | 0.3260 | 0.6234 | 0.4180 | 0.8639 | **0.3148** |
| TCGA-SKCM | 1.0673 | 0.7677 | 0.7316 | 0.6805 | 0.8009 | **0.6499** |
| TCGA-STAD | 0.9864 | 1.0113 | 0.8287 | 0.8560 | **0.8117** | 0.9020 |
| TCGA-THCA | 0.8515 | 0.3891 | 0.4270 | 0.4057 | 0.5880 | **0.3346** |
| TCGA-UCEC | 0.9223 | 0.74911 | 0.7165 | 0.7324 | 0.7587 | **0.7108** |

**Supplementary Table 2.** Mean absolute error (MAE) obtained per model and per cancer type in the five-fold cross-validation joining all test sets on TCGA samples. Best results are highlighted in bold.

| **TCGA Project** | **PloiViT** | | **XGBoost** | | **PloiViT+XGBoost** | |
|---|---|---|---|---|---|---|
| | ResNet | CTrans | ResNet | CTrans | ResNet | CTrans |
| TCGA-BLCA | 0.8968 | 0.6604 | 0.7261 | 0.6884 | 0.6170 | **0.5622** |
| TCGA-BRCA | 1.0633 | 0.7294 | 0.7409 | 0.7184 | 0.8142 | **0.6842** |
| TCGA-GBM | 0.7837 | 0.6212 | 0.6163 | **0.5631** | 0.6743 | 0.5684 |
| TCGA-HNSC | 0.8520 | 0.8644 | **0.6653** | 0.6717 | 0.6721 | 0.7034 |
| TCGA-KICH | 0.8853 | **0.5432** | 0.7047 | 0.6185 | 0.7760 | 0.5555 |
| TCGA-KIRC | 0.7453 | **0.4275** | 0.5831 | 0.5389 | 0.7393 | 0.4562 |
| TCGA-KIRP | 0.7393 | 0.4135 | 0.5073 | 0.3665 | 0.5732 | **0.3427** |
| TCGA-LIHC | 0.7362 | 0.6305 | 0.6886 | 0.6830 | 0.6685 | **0.6281** |
| TCGA-LUAD | 0.9860 | 0.8226 | 0.8756 | 0.8581 | 0.8788 | **0.7979** |
| TCGA-LUSC | 0.8336 | 0.6326 | 0.5440 | **0.5358** | 0.6057 | 0.5448 |
| TCGA-PRAD | 1.023 | **0.2423** | 0.6102 | 0.3999 | 0.7826 | 0.2697 |
| TCGA-SKCM | 0.8730 | 0.6174 | 0.6474 | 0.5630 | 0.6914 | **0.5352** |
| TCGA-STAD | 0.8178 | 0.8270 | 0.6773 | 0.6905 | **0.6684** | 0.7304 |
| TCGA-THCA | 0.7118 | 0.2917 | 0.3976 | 0.3573 | 0.5070 | **0.2495** |
| TCGA-UCEC | 0.7512 | **0.6064** | 0.6549 | 0.6595 | 0.6777 | 0.6262 |

**Supplementary Table 3.** Mean absolute percentage error (MAPE) obtained per model and per cancer type in the five-fold cross-validation joining all test sets on TCGA samples. Best results are highlighted in bold.

| TCGA Project | PloiViT | | XGBoost | | PloiViT+XGBoost | |
|---|---|---|---|---|---|---|
| | ResNet | CTrans | ResNet | CTrans | ResNet | CTrans |
| TCGA-BLCA | 0.3540 | 0.2042 | 0.6777 | 0.2370 | 0.2436 | **0.2039** |
| TCGA-BRCA | 0.4447 | 0.2917 | 0.2723 | 0.2724 | 0.3292 | **0.2687** |
| TCGA-GBM | 0.3693 | 0.2784 | 0.2848 | 0.2561 | 0.3179 | **0.2561** |
| TCGA-HNSC | 0.3248 | 0.3190 | **0.2290** | 0.2394 | 0.2490 | 0.2581 |
| TCGA-KICH | 0.5049 | **0.2728** | 0.3842 | 0.3295 | 0.4364 | 0.2898 |
| TCGA-KIRC | 0.3758 | **0.1833** | 0.2753 | 0.2516 | 0.3180 | 0.2034 |
| TCGA-KIRP | 0.3458 | 0.1734 | 0.2259 | 0.1532 | 0.2666 | **0.1414** |
| TCGA-LIHC | 0.3060 | 0.2555 | 0.2561 | 0.2585 | 0.2637 | **0.2442** |
| TCGA-LUAD | 0.3323 | 0.2765 | 0.2643 | 0.2592 | 0.2813 | **0.2545** |
| TCGA-LUSC | 0.3350 | 0.2438 | **0.2116** | 0.2146 | 0.2450 | 0.2158 |
| TCGA-PRAD | 0.5342 | **0.1260** | 0.3179 | 0.2081 | 0.4086 | 0.1407 |
| TCGA-SKCM | 0.3582 | 0.2289 | 0.2394 | 0.2118 | 0.2770 | **0.2036** |
| TCGA-STAD | 0.3357 | 0.3256 | **0.2507** | 0.2610 | 0.2654 | 0.28371 |
| TCGA-THCA | 0.3593 | 0.1471 | 0.2005 | 0.1806 | 0.2557 | **0.1261** |
| TCGA-UCEC | 0.3452 | **0.2474** | 0.2667 | 0.2688 | 0.2964 | 0.256 |

**Supplementary Table 4.** Root mean squared error (RMSE) obtained per model and per cancer type in the TCGA hold-out test set. Best results are highlighted in bold.

| TCGA Project | PloiViT | | XGBoost | | PloiViT+XGBoost | |
|---|---|---|---|---|---|---|
| | ResNet | CTrans | ResNet | CTrans | ResNet | CTrans |
| TCGA-BLCA | 1.1677 | **0.1572** | 0.7154 | 0.6601 | 0.9388 | 0.2514 |
| TCGA-BRCA | 0.7477 | 0.7898 | 0.5210 | **0.4696** | 0.5545 | 0.5488 |
| TCGA-GBM | **0.6205** | 0.9569 | 0.6509 | 0.6549 | 0.6177 | 0.7036 |
| TCGA-HNSC | 0.7504 | 1.5197 | 0.8010 | 0.7874 | **0.7335** | 1.1503 |
| TCGA-KICH | 0.9753 | 0.6240 | 0.6916 | 0.8485 | 0.8280 | **0.6028** |
| TCGA-KIRC | 0.6832 | 0.3244 | 0.5232 | 0.5779 | 0.5856 | **0.13113** |
| TCGA-KIRP | 0.5720 | 0.4848 | 0.4107 | 0.4323 | 0.4729 | **0.1126** |
| TCGA-LIHC | **0.6336** | 1.0575 | 0.7494 | 0.6892 | 0.6871 | 0.8021 |
| TCGA-LUAD | **1.008** | 1.5387 | 1.1337 | 1.1924 | 1.0477 | 1.3489 |
| TCGA-LUSC | 1.0437 | 0.8017 | 0.7152 | 0.7208 | 0.8461 | **0.6855** |
| TCGA-PRAD | 1.0572 | 0.2838 | 0.5963 | 0.5823 | 0.8235 | **0.1502** |
| TCGA-SKCM | 0.4968 | 0.5557 | 0.1743 | 0.3187 | 0.3355 | **0.1184** |
| TCGA-STAD | **0.2238** | 1.3485 | 0.7682 | 0.5237 | 0.2722 | 0.9361 |
| TCGA-THCA | 0.7093 | 0.2763 | 0.4430 | 0.5823 | 0.5738 | **0.1535** |
| TCGA-UCEC | 0.6762 | 0.7735 | 0.5066 | **0.4400** | 0.5350 | 0.5011 |

**Supplementary Table 5.** Mean absolute error (MAE) obtained per model and per cancer type in the TCGA hold-out test set. Best results are highlighted in bold.

| TCGA Project | PloiViT | | XGBoost | | PloiViT+XGBoost | |
|---|---|---|---|---|---|---|
| | ResNet | CTrans | ResNet | CTrans | ResNet | CTrans |
| TCGA-BLCA | 1.1603 | **0.1568** | 0.6891 | 0.6591 | 0.9247 | 0.2511 |
| TCGA-BRCA | 0.6258 | 0.6325 | 0.4498 | **0.4260** | 0.5039 | 0.4366 |
| TCGA-GBM | 0.4831 | 0.7195 | 0.4881 | 0.6024 | 0.4571 | **0.3793** |
| TCGA-HNSC | **0.4495** | 1.5020 | 0.7655 | 0.7498 | 0.5976 | 1.1259 |
| TCGA-KICH | 0.9150 | **0.3419** | 0.6551 | 0.8245 | 0.7851 | 0.4972 |
| TCGA-KIRC | 0.6515 | 0.3223 | 0.5081 | 0.5772 | 0.5798 | **0.1274** |
| TCGA-KIRP | 0.4811 | 0.4721 | 0.3716 | 0.4173 | 0.4177 | **0.1033** |
| TCGA-LIHC | **0.5195** | 0.8300 | 0.6773 | 0.6474 | 0.5984 | 0.6174 |
| TCGA-LUAD | 0.7688 | 1.1983 | 0.8097 | 0.8410 | **0.7353** | 0.9687 |
| TCGA-LUSC | 0.8979 | **0.5779** | 0.6919 | 0.7102 | 0.7567 | 0.6112 |
| TCGA-PRAD | 1.0386 | 0.2824 | 0.5783 | 0.5815 | 0.8085 | **0.1495** |
| TCGA-SKCM | 0.4968 | 0.5557 | 0.1743 | 0.3187 | 0.3355 | **0.1184** |
| TCGA-STAD | **0.2238** | 1.3485 | 0.7682 | 0.5237 | 0.2722 | 0.9361 |
| TCGA-THCA | 0.6894 | 0.2762 | 0.4424 | 0.5819 | 0.5659 | **0.1528** |
| TCGA-UCEC | 0.5873 | 0.6156 | 0.4418 | 0.4176 | 0.5031 | **0.3618** |

**Supplementary Table 6.** Mean absolute percentage error (MAPE) obtained per model and per cancer type in the TCGA hold-out test set. Best results are highlighted in bold.

| TCGA Project | PloiViT | | XGBoost | | PloiViT+XGBoost | |
|---|---|---|---|---|---|---|
| | ResNet | CTrans | ResNet | CTrans | ResNet | CTrans |
| TCGA-BLCA | 0.6212 | **0.0840** | 0.3684 | 0.3533 | 0.4948 | 0.1346 |
| TCGA-BRCA | 0.2826 | 0.2273 | 0.1950 | 0.1876 | 0.2275 | **0.1642** |
| TCGA-GBM | 0.1850 | 0.2601 | 0.1768 | 0.2405 | 0.1671 | **0.1143** |
| TCGA-HNSC | **0.1248** | 0.4555 | 0.2296 | 0.2247 | 0.1740 | 0.3401 |
| TCGA-KICH | 0.5185 | 0.1345 | 0.3750 | 0.4730 | 0.4468 | 0.2607 |
| TCGA-KIRC | 0.3287 | 0.1627 | 0.2578 | 0.2923 | 0.2932 | **0.0647** |
| TCGA-KIRP | 0.2273 | 0.2196 | 0.1769 | 0.1990 | 0.1982 | **0.0481** |
| TCGA-LIHC | **0.1885** | 0.2802 | 0.2559 | 0.2597 | 0.2222 | 0.2072 |
| TCGA-LUAD | 0.2112 | 0.3221 | 0.2084 | 0.2123 | **0.1909** | 0.2450 |
| TCGA-LUSC | 0.4586 | **0.2094** | 0.3375 | 0.3352 | 0.3857 | 0.2550 |
| TCGA-PRAD | 0.5353 | 0.1455 | 0.2981 | 0.2997 | 0.4167 | **0.0771** |
| TCGA-SKCM | 0.2214 | 0.2476 | 0.0776 | 0.1420 | 0.1495 | **0.0528** |
| TCGA-STAD | **0.0731** | 0.4409 | 0.2512 | 0.1712 | 0.0890 | 0.3061 |
| TCGA-THCA | 0.3572 | 0.1431 | 0.2292 | 0.3015 | 0.2932 | **0.0791** |
| TCGA-UCEC | 0.2738 | 0.2336 | 0.1849 | 0.1778 | 0.2244 | **0.1342** |

**Supplementary Table 7.** Mean absolute error (MAE) obtained per model and per cancer type in the CPTAC cohort. Best results are highlighted in bold.

| CPTAC Project | PloiViT | | XGBoost | | PloiViT+XGBoost | |
|---|---|---|---|---|---|---|
| | ResNet | CTrans | ResNet | CTrans | ResNet | CTrans |
| CPTAC-GBM | 0.6306 | **0.5499** | 0.7133 | 0.6111 | 0.6702 | 0.5775 |
| CPTAC-LUAD | 1.063 | **0.9216** | 1.0726 | 1.0226 | 1.0671 | 0.9691 |

**Supplementary Table 8.** Mean absolute percentage error (MAPE) obtained per model and per cancer type in the CPTAC cohort. Best results are highlighted in bold.

| CPTAC Project | PloiViT | | XGBoost | | PloiViT+XGBoost | |
|---|---|---|---|---|---|---|
| | ResNet | CTrans | ResNet | CTrans | ResNet | CTrans |
| CPTAC-GBM | 0.2352 | **0.1904** | 0.2876 | 0.2236 | 0.2606 | 0.2056 |
| CPTAC-LUAD | 0.2859 | **0.2510** | 0.2855 | 0.2784 | 0.2854 | 0.2635 |

**Supplementary Table 9.** Mean absolute error (MAE) obtained per model and slide in the spatial glioblastoma cohort. Given the low performance of the XGBoost model, we did not compute the performance of the PloiViT+XGBoost model. Best results are highlighted in bold.

| Slide ID | PloiViT | | XGBoost | |
|---|---|---|---|---|
| | ResNet | CTrans | ResNet | CTrans |
| 269_T | 0.1018 | **0.0446** | 0.6255 | 0.7731 |
| 270_T | 0.1445 | **0.0936** | 0.6900 | 0.7111 |
| 296_T | **0.1621** | 0.1792 | 0.5588 | 0.7640 |
| 275_T | 0.0663 | **0.0221** | 0.6694 | 0.5213 |
| 248_T | 0.0306 | **0.0125** | 0.5939 | 0.6429 |
| 334_T | 0.2575 | **0.0438** | 0.6488 | 0.5899 |
| 268_T | 0.1156 | **0.0603** | 0.6428 | 0.7188 |
| 255_T | **0.1228** | 0.1841 | 0.6007 | 0.6275 |
| 259_T | **0.0408** | 0.0491 | 0.6208 | 0.7112 |
| 266_T | **0.0293** | 0.1131 | 0.7095 | 0.6877 |
| 242_T | 0.1384 | **0.0705** | 0.6287 | 0.9284 |
| 262_T | 0.1258 | **0.0990** | 0.6782 | 0.6160 |
| 260_T | 0.0659 | **0.0231** | 0.5568 | 0.7639 |
| 243_T | **0.0292** | 0.0441 | 0.5827 | 0.7643 |
| 265_T | **0.2038** | 0.2092 | 0.5858 | 0.6009 |
| 304_T | **0.1173** | 0.1264 | 0.6174 | 0.6399 |
| Mean | 0.1095 | **0.0859** | 0.6256 | 0.6913 |
| STD | 0.0631 | **0.0597** | 0.0440 | 0.0943 |

**Supplementary Table 10.** Mean absolute percentage error (MAPE) obtained per model and slide in the spatial glioblastoma cohort. Given the low performance of the XGBoost model, we did not compute the performance of the PloiViT+XGBoost model. Best results are highlighted in bold.

| **Slide ID** | **PloiViT** | | **XGBoost** | |
|---|---|---|---|---|
| | ResNet | CTrans | ResNet | CTrans |
| 269_T | 0.0509 | **0.0223** | 0.3126 | 0.3864 |
| 270_T | 0.0721 | **0.0467** | 0.3445 | 0.3551 |
| 296_T | **0.0810** | 0.0896 | 0.2794 | 0.3820 |
| 275_T | 0.0331 | **0.0110** | 0.3345 | 0.2605 |
| 248_T | 0.0152 | **0.0062** | 0.2968 | 0.3213 |
| 334_T | 0.1288 | **0.0219** | 0.3246 | 0.2951 |
| 268_T | 0.0577 | **0.0301** | 0.3211 | 0.3590 |
| 255_T | **0.0614** | 0.0920 | 0.3003 | 0.3137 |
| 259_T | **0.0204** | 0.0245 | 0.3098 | 0.3549 |
| 266_T | **0.0146** | 0.0565 | 0.3546 | 0.3437 |
| 242_T | 0.0691 | **0.0352** | 0.3140 | 0.4637 |
| 262_T | 0.0628 | **0.0494** | 0.3388 | 0.3078 |
| 260_T | 0.0330 | **0.0115** | 0.2792 | 0.3830 |
| 243_T | **0.0146** | 0.0220 | 0.2913 | 0.3820 |
| 265_T | **0.1018** | 0.1045 | 0.2925 | 0.3000 |
| 304_T | **0.0585** | 0.0631 | 0.3083 | 0.3195 |
| Mean | 0.0547 | **0.0429** | 0.3126 | 0.3455 |
| STD | 0.0315 | **0.0298** | 0.0218 | 0.0471 |

**Supplementary Table 11.** Number of Whole Slide Images and unique number of patients used from each cancer type in TCGA.

| **TCGA project** | **number WSIs** | **number patients** |
|---|---|---|
| TCGA-BLCA | 23 | 22 |
| TCGA-BRCA | 90 | 88 |
| TCGA-GBM | 155 | 39 |
| TCGA-HNSC | 29 | 29 |
| TCGA-KICH | 43 | 43 |
| TCGA-KIRC | 34 | 34 |
| TCGA-KIRP | 94 | 32 |
| TCGA-LIHC | 51 | 51 |
| TCGA-LUAD | 126 | 37 |
| TCGA-LUSC | 179 | 47 |
| TCGA-PRAD | 55 | 19 |
| TCGA-SKCM | 42 | 36 |
| TCGA-STAD | 40 | 34 |
| TCGA-THCA | 49 | 48 |
| TCGA-UCEC | 48 | 43 |
| total | 1058 | 602 |

**Supplementary Table 12.** Number of Whole Slide Images and unique number of patients used from each cancer type in CPTAC.

| **CPTAC project** | **number WSIs** | **number patients** |
|---|---|---|
| CPTAC-GBM | 464 | 100 |
| CPTAC-LUAD | 462 | 95 |
| total | 926 | 195 |

**Supplementary Table 13.** Number of Whole Slide Images and unique number of patients used from each cancer type in PBTA.

| Cancer type | number WSIs | number patients |
|---|---|---|
| Low-grade glioma/astrocytoma (LGG) | 84 | 45 |
| High-grade glioma/astrocytoma (HGG) | 36 | 11 |
| Ganglioglioma | 2 | 1 |
| Other | 3 | 1 |
| total | 125 | 58 |
